# Loss of Sarm1 Mitigates Axonal Degeneration and Promotes Neuronal Repair After Ischemic Stroke

**DOI:** 10.1101/2025.02.20.639171

**Authors:** Jack T. Wang, Jennifer An, Brian Toh, Kasra Yousefpour, Yutaro Komuro, Marlesa I. Godoy, Jennifer Putman, S. Thomas Carmichael, Robert Damoiseaux, Jason D. Hinman

## Abstract

Axonal degeneration is a core feature of ischemic brain injury that limits functional recovery^1^. The pro-degenerative molecule Sarm1 is required for Wallerian axon degeneration after traumatic and chemotoxic nerve injuries^2^, however it is unclear if a similar mechanism mediates axonal degradation after ischemic injury. Here we show that loss of Sarm1 results in profound attenuation of axonal degeneration after focal ischemia to the subcortical white matter as well as to the cortex. Moreover, absence of Sarm1 significantly promotes the survival of neurons remote from but connected to the infarct after ischemic injuries to the subcortical white matter as well as to the cortex. Notably, loss of Sarm1 also significantly ameliorates early and late motor as well as cognitive deficits following white matter stroke. To further understand the mechanism of *Sarm1-/-* mediated neuronal protection, we performed differential gene expression analyses of wildtype and *Sarm1-/-* stroke-injured neurons and found that the loss of Sarm1 activates a pro-growth molecular program that promotes gene expression programs involved in axonogenesis and synaptogenesis after white matter ischemia. Using a functional genomics approach to recapitulate such a molecular program in *Sarm1-/-* neurons, we identify molecular compounds sufficient to enhance cortical neurite outgrowth *in vitro*, and all of which elicit a conserved epigenetic signature promoting axonogenesis. These results indicate that Sarm1 promotes axonal degeneration and concurrently inhibits an axonal reparative program encoded at the level of the epigenome that can be modulated pharmacologically. Our findings thus reveal a novel role for Sarm1 as a crucial regulator of both axonal degeneration and axonal repair after ischemic stroke.

**SIGNIFICANCE STATEMENT:** Axon degeneration is a pivotal event following ischemic stroke, however the mechanism of white matter loss in stroke is unknown. We demonstrate that the pro-degenerative molecule Sarm1 is required for axonal and neuronal degeneration after ischemic injuries, improves functional motor and cognitive recovery after stroke, and that loss of Sarm1 surprisingly induces the activation of a reparative program driving new axon and synapse formation. Using functional genomics, we uncover molecular candidates that phenocopy this pro-growth molecular signature in *Sarm1-/-* neurons and show that these compounds are sufficient to promote de novo axonal growth via an epigenetic mechanism. Our results thus reveal a novel role for Sarm1 as a regulator of both axonal degeneration and axonal repair after ischemia, in addition to identifying pharmacologic candidates to promote axonal remodeling in stroke.

## INTRODUCTION

Axonal degeneration is a characteristic event in ischemic stroke^3^, and the degree of axon loss in the various cortical and spinal white matter tracts is an independent determinant of functional recovery for stroke patients^1,4,5^. However, comparatively little is known about the mechanisms mediating axonal degeneration and subsequent white matter remodeling after ischemia.

Ischemic stroke results in spatially and temporally distinct phases of injury to the white matter^6^, the severity of which impacts not only the integrity and connectivity of the white matter, but also the extent of subsequent attempted axonal repair and remodeling^7,8^. This involves first an acute, Wallerian-like phase of degradation that is related to direct, local injury to the tissue, and results in local bioenergy failure and rapid disintegration of the axon via calcium influx and activation of Ca2+ dependent proteases^9^. In mice, ischemia triggers a similar morphologic sequence of axonal degradation as traumatic nerve injuries that involve local axonal swelling and beading, followed by rapid fragmentation of the axolemma^10,11^. Similar morphologic changes in the white matter are also observed radiographically via diffusion tensor imaging in stroke patients^12,13,14,15,16^. This acute phase is then followed by a delayed, secondary phase that damages the surviving but vulnerable axons near the penumbra likely through axon-glial interactions and surrounding inflammation^7^.

The significant morphologic similarities in axonal degeneration in both ischemic and traumatic nerve injuries indicate potentially shared underlying molecular mechanism(s) mediating the process. Thus, potential insight may be gleaned from our existing understanding of Wallerian degeneration caused by other modes of CNS injuries. Prior studies have shown that the pro-degenerative protein Sarm1 (Sterile Alpha and TIR Motif containing 1) is required for axon degeneration in traumatic and chemotoxic injuries. Sarm1 is a conserved protein containing two sterile-alpha motifs (SAM), one mitochondrial association (MT) and one TIR domain with intrinsic NADase activity^17,18,19,20^. Loss of Sarm1 confers protection from axonopathy triggered by traumatic injury, chemotherapeutic agents, and mitochondrial dysfunction^21,22,23,24^. It is thought that Sarm1 triggers axonal degeneration after traumatic injury via its NAD+ glycohydrolase activity, and the resultant decrease in overall bioenergy content, imbalance in intracellular NAD/NMN ratio, or the generation of toxic NAD+ metabolites ultimately leads to increased Ca2+ influx and subsequent axonal cytoskeletal breakdown by Ca2+ dependent proteases^9,25,26,27^.

Whether a similar Sarm1-dependent mechanism regulates axonal degeneration after ischemia is unclear. No direct link between Sarm1 activity and axonal degradation after ischemia has been established to date, and the effect of delaying Wallerian degeneration on subsequent attempted CNS neurite regrowth and repair after stroke is not known. In this study, we use well characterized and robust murine models of subcortical white matter stroke^6,28^ as well as cortical stroke^29,30^ to investigate the effects of Sarm1 loss on axonal as well as neuronal survival after ischemic injury. We show that loss of Sarm1 results in robust axonal as well as neuronal somal protection following injury in two different *in vivo* models of ischemic stroke. To better understand the mechanism of *Sarm1-/-* mediated axonal and neuronal protection, we compare the transcriptional signatures in stroke-injured cortical neurons and surprisingly reveal a novel pro-growth molecular program from loss of Sarm1. Using a functional genomics approach, we further identify molecular candidates that replicate the protective molecular signature in *Sarm1-/-* neurons and are functionally sufficient to promote axonal outgrowth. Our study thus provides insight into the mechanism of Sarm1-mediated axonal and neuronal degeneration in stroke, and identifies novel therapeutic candidates to potentially ameliorate axon degeneration and secondary neurodegeneration, as well as to promote axonal repair and remodeling after ischemia.

## RESULTS

### Loss of Sarm1 attenuates axonal degeneration and results in persistent survival in stroke-injured neurons

We first sought to determine the mechanism of axonal loss in ischemic injuries. As axonal degeneration following ischemia shares morphological similarities with Sarm1-dependent axonal loss after traumatic nerve injuries^17,18,19^, we hypothesized that Sarm1 activity is also required for axonal degeneration after ischemic stroke. We employed an *in vivo* model of ischemic white matter stroke^6,10,28^ and compared axonal survival between *Sarm1-*null (*Sarm1-/-*) and wildtype (WT) mice. Focal ischemic injury to the white matter is achieved via stereotactic delivery of an irreversible eNOS inhibitor L-N^5^-(1-Iminoethyl) ornithine, Dihydrochloride (L-Nio) to antero-lateral region of the corpus callosum to generate a local vasoconstrictive and ischemic environment in the white matter. A retrograde fluorescent tracer (fluororuby; FR) concomitantly delivered at the same time as L-Nio-induced ischemic injury aids in labeling and identifying stroke-injured neurons and axonal projections^31^ (Fig. 1A). Mean neurite fluorescence of FR+ callosal axon projections distal to the subcortical stroke are preserved throughout the corpus callosum in *Sarm1-/-* mice while they are significantly attenuated in WT mice at 7 days (mean neurite fluorescence 20.78 in WT vs 255.70 in *Sarm1-/-* mice; *p*<0.0001) and at 28 days (mean 10.59 in WT vs 79.53 in *Sarm1-/-* mice; *p*<0.0001) after ischemic stroke (Fig. 1B-E). In ultrastructural electron microscopy samples from the midline corpus callosum, *Sarm1-/-* mice demonstrate relative preservation of axons with increased numbers of intact and organized peri- and endoneural structures, with a trend towards increased myelinated axons per ultrastructural field of view during the first week after stroke (3d: 66.9±20.8 vs. 74.3±11.7; 7d: 69.9±23.3 vs. 114.6±21.4; *p*=0.35 for interaction by two-way ANOVA; Fig. 1B, F). Thus, loss of Sarm1 activity robustly confers long-distance and long-lasting (up to 28 days tested) axonal preservation after ischemic stroke.

**Figure 1.**
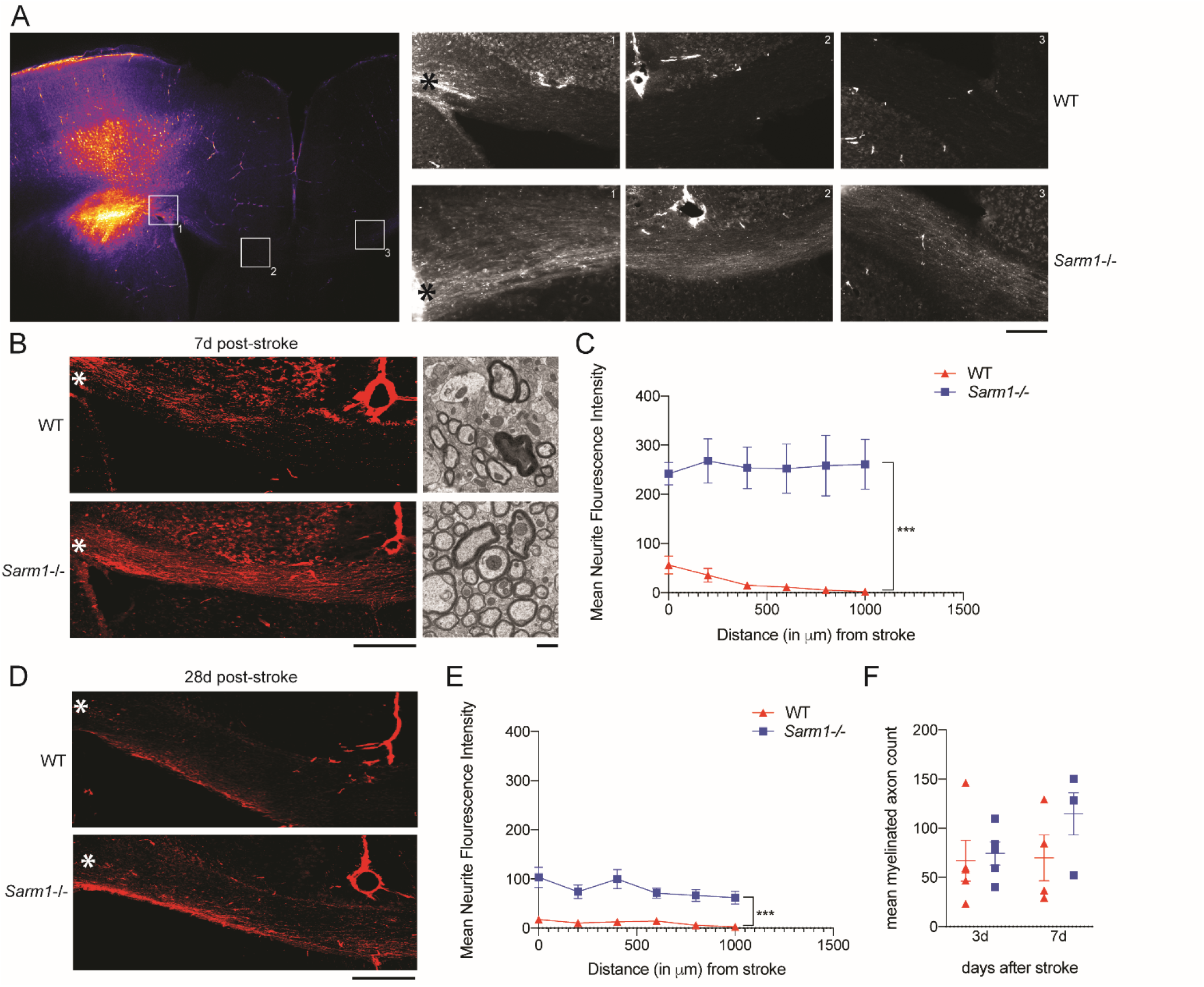
Loss of Sarm1 results in persistent axonal survival after white matter stroke. Focal white matter stroke with concurrent retrograde tracer administration (fluororuby, FR) targets subcortical white matter (right, A) and differentially labels transcallosal distal axonal projections 7 days after stroke between wild-type (WT) (upper left, A) and *Sarm1-/-* mice (lower left, A). Numbered white boxes denote corresponding topographic location along the corpus callosum. Representative confocal microscopy (right, B) and electron microscopy images (left, B) from the midline corpus callosum (B) of WT and *Sarm1-/-* brain tissues at 7 days after white matter stroke. Mean neurite FR intensities per unit distance ipsilateral to the stroke lesion at 7 days post-stroke (\*\*\**p*<0.001 by Student t-test, *n*=9 per genotype) (C). Representative confocal microscopy images of WT and *Sarm1-/-* brain tissues at 28 days after white matter stroke (D). Mean neurite FR intensities per unit distance ipsilateral to the stroke lesion at 28 days post-stroke (\*\*\**p*<0.001 by Student t-test, *n*=9 per genotype). Quantification of myelinated axons/FOV in WT and *Sarm1-/-* axons per midline callosal cross section (*n*=4/genotype/timepoint) (F). Asterisks denote the lateral edge of stroke lesion. Error bars=SEM. Scale bars = 100 μm (A); 80 μm (B, right), 1 μm (B, left).

As focal, subcortical ischemic stroke is associated with selective neuronal degeneration distally in the cortex^32^, we investigated how delaying axonal degeneration after stroke injury to the subcortical white matter may impact the survival of cortical neurons distant from but anatomically connected to the stroke lesion. We performed tissue clearing of stroke-injured brains using the previously described uDISCO technique^33^ to resolve FR+ stroke-injured neurons between WT and *Sarm1-/-* mice. We confirmed that the FR+ cells were predominantly neuronal based on NeuN+ co-localization, thus demonstrating specific FR+ labeling of stroke-injured neurons (Fig 2B). Using 3D volumetric analysis of the uDisco-cleared tissue sections, we identified a significant increase in the density of stroke-injured FR+ cortical neurons in *Sarm1-/-* animals compared to WT mice at 7 days after ischemic white matter stroke (mean cell density in WT vs. *Sarm1-/-* 49.2±3.48 cells vs. 81.8±11.5 cells per 3x10^6^ μm^3^ of cortical tissue, respectively; *p*=0.043 by unpaired t-test) (Figure 2A-C). To further determine whether the *Sarm1-/-* dependent protective effect on remote but connected stroke-injured neurons is conserved across different stroke injuries with varying severity and infarct locations, we also compared the survival of subcortical thalamic neurons following cortical ischemia via permanent distal middle cerebral artery occlusion (dMCAO) between wildtype and *Sarm1-/-* animals. We found that loss of Sarm1 similarly improved the survival of labelled (via AAV-DJ-EGFP transduction) subcortical thalamic neurons 7 days after a cortical stroke from dMCAO, as evidenced by the significantly greater number of NeuN+ cells in the ipsilateral thalamus of *Sarm1-/-* mice than that of WT mice (mean thalamic NeuN+ cells 31.25 ± 4.07 in wildtype vs 46.50 ± 5.24 in *Sarm1-/-* mice; *p*=0.002) (Fig 2D-E). Thus, the absence of Sarm1 significantly ameliorated stroke-induced neuronal degeneration in two different *in vivo* stroke models.

**Figure 2.**
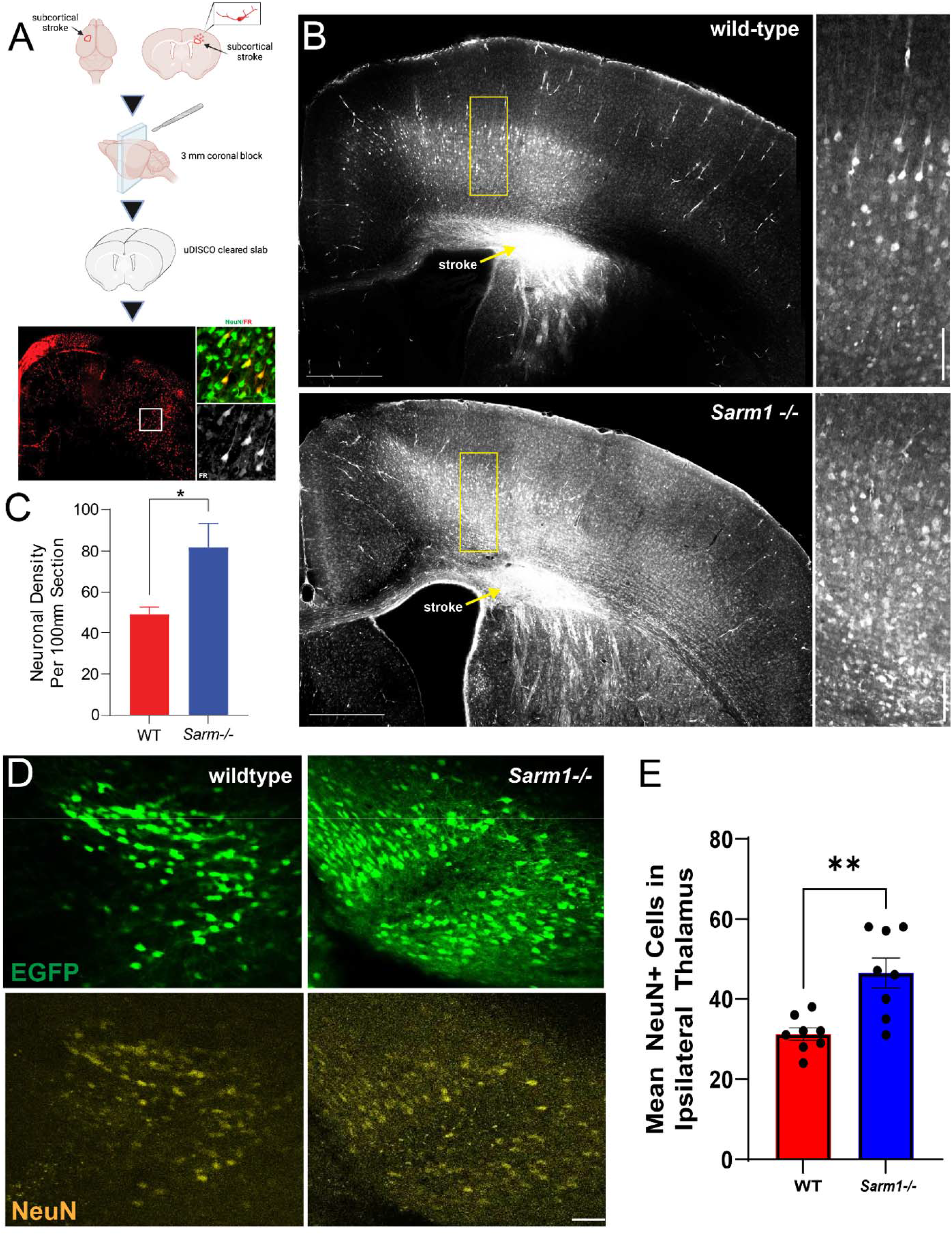
Loss of Sarm1 promotes neuronal survival after ischemic stroke. uDISCO clearing of brain tissue slabs (3 mm) in WT and *Sarm1-/-* mice at 7 days after subcortical white matter stroke results in volumetric visualization of fluororuby (FR)+ cortical neurons that also label for NeuN (A). Representative optical slices from reconstructed 3D image from WT (upper) and *Sarm1-/-* (lower) mice demonstrates retrograde labeling of stroke-injured surviving neuronal cells in overlying sensorimotor cortex (B). Density of surviving FR+ cortical neurons ipsilateral to the infarct as measured and quantified by using semi-automated 3D cell counter plugin in Image J (\**p*=0.04 by Student’s t-test, n=4/genotype) (C). Cortical as well as subcortical thalamic neurons are fluorescently labelled by injection of AAV-hsyn-EGFP reporter construct to the cortex, with immunohistochemical staining for NeuN+ and topographic identification of the thalamic structure to confirm labeling of thalamic neurons ipsilateral to the infarct at 7 days after cortical stroke via permanent distal middle cerebral artery occlusion (dMCAO) (D). Mean number of NeuN+ thalamic neurons between wildtype and *Sarm1-/-*mice is quantified at 7 days after cortical dMCAO stroke (\*\**p*=0.01 by Student’s t-test, n=8/genotype) (E). Error bars = SEM. Scale bars = 500um (B, left); 100um (B, right); and 100 um (C).

### Loss of Sarm1 improves functional motor and cognitive recovery after ischemic stroke

To determine whether loss of Sarm1, and thus attenuation of axonal as well as neuronal degeneration, leads to differences in behavioral and functional outcomes after ischemic stroke, we compared the motor and cognitive functions between wildtype and *Sarm1-/-* mice at up to 30 days after ischemic stroke. We observed that on the grid-walking task, *Sarm1 /* mice showed significantly fewer ipsilateral forelimb foot faults than WT mice across the recovery period (adjusted *p*=0.006, two-way repeated-measures ANOVA; **Figure 3A**). Similarly, on the cylinder task, *Sarm1 /* mice exhibited significantly reduced ipsilateral forepaw-use bias relative to WT mice (adjusted *p*=0.03, two-way repeated-measures ANOVA; **Figure 3B**), reflecting more symmetric, and thus improved, forelimb use after stroke. Finally, using a fear conditioning paradigm to assess associative learning and memory, we found that the percent change in freezing behavior between training and retrieval differed significantly between genotypes (mean delta freezing WT vs. Sarm1-/-, 3.14 ± 2.19 vs. 12.2 ± 2.02*, p=*0.008 by unpaired t-test; **Figure 3C**), suggesting that Sarm1 deletion also preserves cognitive function after ischemic stroke. Thus, loss of Sarm1 significantly ameliorated the motor and cognitive impairment typically observed in this subcortical ischemic stroke model.

**Figure 3.**
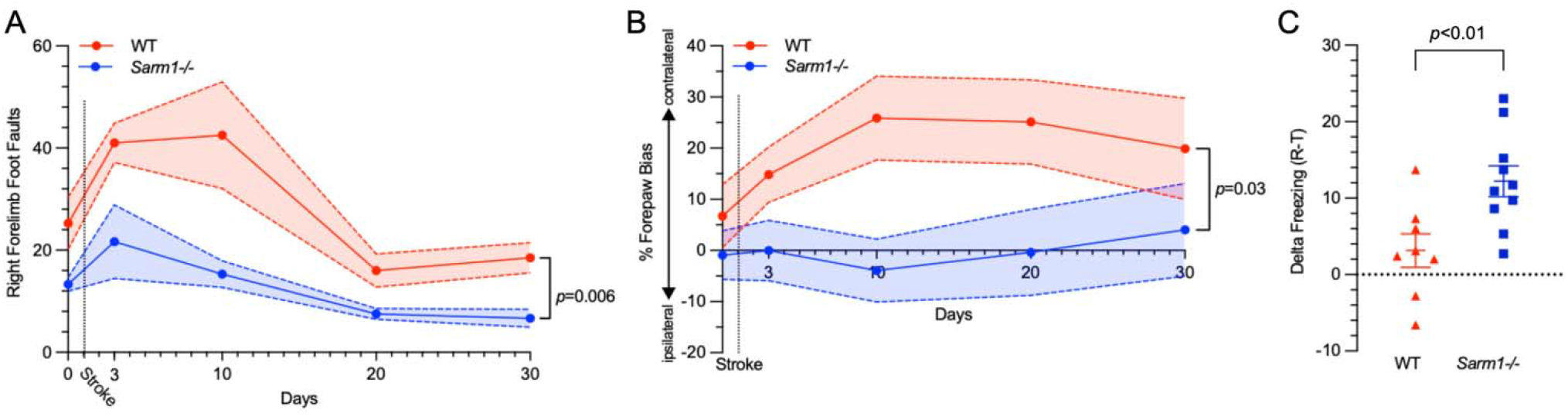
Loss of Sarm1 mitigates post-stroke motor and cognitive deficits after stroke. Wild-type (*n*=8) or *Sarm1*-/- animals (*n=*10) underwent baseline (Day 0) behavioral assessment of motor function on grid-walking (A) and cylinder tasks (B) followed by subcortical stroke (Day 1). Serial assessments of motor function were performed on Days 3, 10, 20 and 30 followed by fear conditioning training and retrieval (C). Ipsilateral (right) forelimb foot faults on grid-walking were significantly different between WT and *Sarm1*-/- animals (adjusted *p*=0.006 by two-way repeated measures ANOVA) (A). Ipsilateral forepaw bias on cylinder task was significantly different between WT and *Sarm1*-/- animals (adjusted *p*=0.03 by two-way repeated measures ANOVA) (B). Percent change in freezing between training and retrieval was significantly different between WT and *Sarm1*-/- animals (*p*<0.01 by unpaired t-test) (C). Error bars = SEM.

### Activation of pro-growth transcriptional pathways in surviving stroke-injured neurons

We further reasoned that this post-stroke enhancement of neuronal survival in the axonally protective *Sarm1-/-* mice occurs via transcriptional regulation. To determine the effect of *Sarm1* deletion on gene expression by the cortical neurons after ischemic injury, we utilized MACS-FACS-seq to transcriptionally profile stroke-injured cortical neurons of *Sarm1-/-* and WT mice at 7 days after white matter ischemic stroke (Fig. 4A). Using a FR+/NCAM+ MACS-FACS isolation strategy, we sorted similar numbers of cells in WT and *Sarm1-/-* mice (5,788±3,171 vs. 4,207±1,743 FR+/NCAM+ cells; *p*=0.68 by unpaired t-test; and see also Fig. S1) that enrich for deep cortical layer neurons without contamination of other non-neuronal cell types (neuronal enrichment *p*=0.00042 by one-way ANOVA, F_(4,11)_=12.68, Fig. 4B). Analysis of differentially expressed genes (DEG) (FDR<0.05) between *Sarm1*-/-and WT stroke-injured cortical neurons identified a unique set of 891 changing genes (621 up-regulated, 270 down-regulated) (Fig. 4C; and see also Fig. S2 and Table S1). The top 150 up-regulated genes in *Sarm1-*/- neurons are functionally correlated with those implicated in neuronal differentiation, axonal outgrowth, synaptic transmission, calcium signaling, and neuronal metabolism^34^ (see also Table S2). Notably, genes previously implicated in the defined Sarm1 signaling pathways^35,36^ were not significantly altered in *Sarm1*-/- mice compared to WT mice (Fig. 4D), suggesting that the transcriptional response of cortical neurons to ischemic injury in the absence of *Sarm1* does not directly involve activation of known Sarm1 pathways.

**Figure 4.**
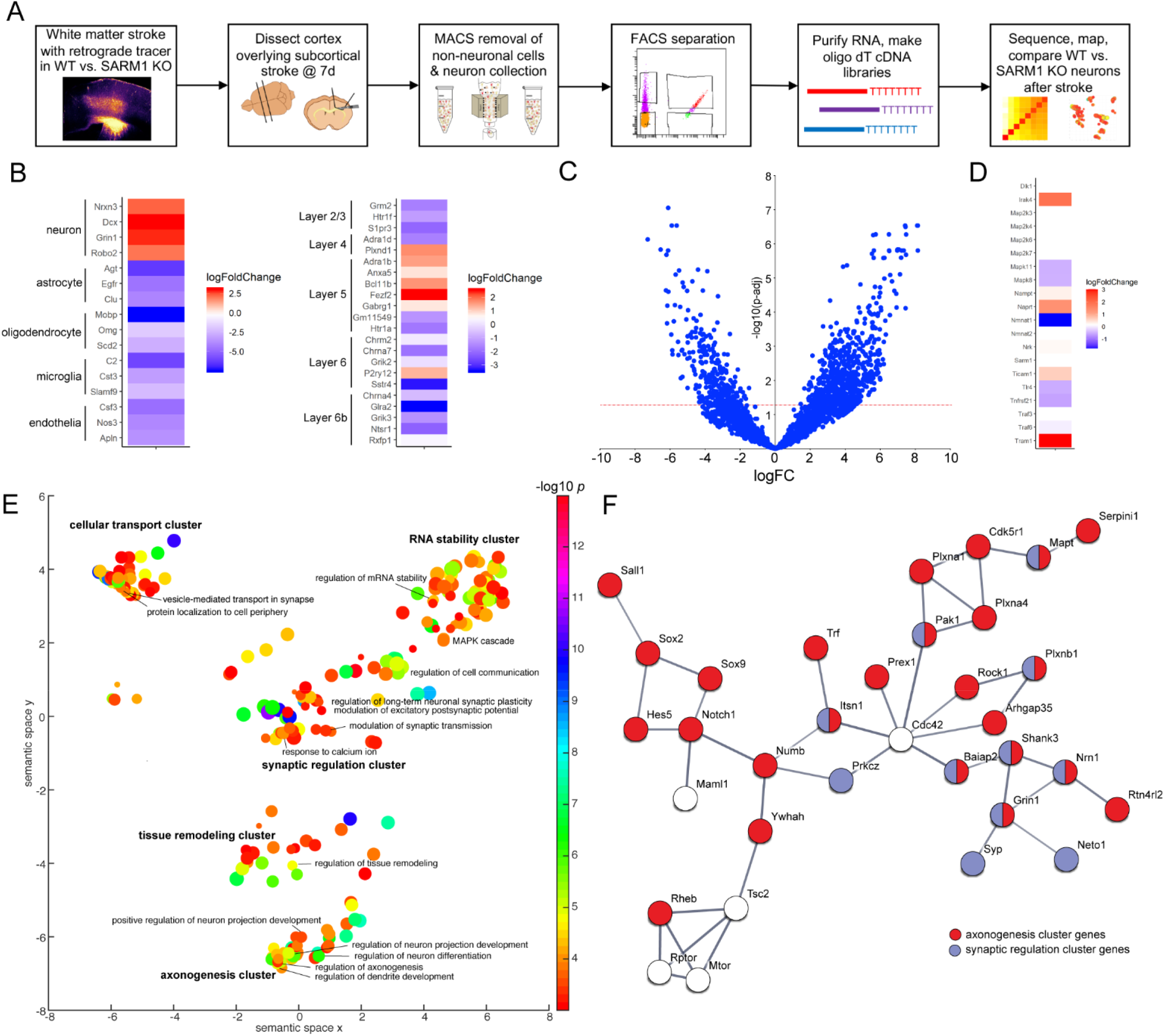
Genetic deletion of *Sarm1* activates axonogenesis and synaptogenesis in stroke-injured cortical neurons. Schematic model for MACS-FACS-seq cell isolation workflow in wild-type (WT) and *Sarm1*-/- mice (A). Quantification of number of *Sarm1-/-* and WT FACS-sorted NCAM+ and fluororuby +/- neurons is further shown in Figure S1. Log fold-change (logFC) expression of cell-type marker (left) and cortical layer (right) genes demonstrating enrichment of deep cortical layer neurons by MACS-FACS-seq (B). Volcano plot of 891 differentially expressed genes (DEGs) (FDR<0.05) between *Sarm1*-/- and WT neurons seven days after subcortical stroke (C). Heat map expression of all DEGs between *Sarm1-/-* and WT neurons is further shown in Figure S2. Heat-map of logFC expression changes in known Sarm1 pathway genes between *Sarm1*-/- and WT neurons after stroke (all FDR>0.05) (D). REVIGO semantic clustering map of gene ontology terms derived from stroke-induced *Sarm1*-/-DEG profile showing five major clusters of activity (bold) (color bar = -log_10_ *p*-value) (E). Expression levels of the gene clusters between *Sarm1-/-* and WT neurons after stroke are further shown in Figure S3. STRING protein-protein interaction network derived from genes in the axonogenesis (red; PPI enrichment *p*=3.88^e-08^) and synaptogenesis (blue; PPI enrichment *p*=8.84^e-08^) clusters (F). Heat map of logFC expression changes of genes within the axonogenesis and synaptogenesis clusters is further shown in Figure S4.

We then utilized REVIGO to organize the gene ontology terms derived from this DEG profile and identified five clusters of molecular programs enriched in stroke-injured cortical neurons in the absence of *Sarm1* (Fig. 4E; and see also Table S3): *cellular transport*; *RNA stability*; *tissue remodeling*, *synaptic regulation*, and *axonogenesis*. Consistent with a predicted impact on retrograde axon-to-cell body signaling in the absence of *Sarm1*^37,38^, the *cellular transport* cluster is enriched for GO terms including protein localization to cell periphery (GO:1990778) and vesicle-mediated transport (GO:0016192), driven in part by strong up-regulation of genes such as *Ezr* (logFC = 3.11) and *Tulp3* (logFC = 3.83) with recognized roles in these processes (Fig S3). Similarly, the *RNA stability* cluster is marked by GO terms and genes implicated in the regulation of RNA stability, a key feature of retrograde axon signaling (GO:0061157 – *Rbm24*; GO:0043488 – *Khsrp*)^39^. Unexpectedly, we also identified two distinct clusters: *axonogenesis* and *synaptic regulation*. This indicates that a pro-growth, regenerative state may be induced in stroke-injured cortical neurons in the absence of *Sarm1*. These clusters feature numerous differentially expressed genes implicated in neuron differentiation (*Bcl11a, Tbr1, Nap1l2*), axonogenesis (*Cdk5r1, Itga4, Serpini1*), and synaptic formation (*Crtc1, Itsn1, Prkcz*) (Table 1; and see also Fig. S3 and Fig S4). Notably, the genes within these regenerative clusters demonstrate a high degree of connectivity with each other and tightly associate within a predicted protein-protein interaction network centering around the recognized pro-growth molecule Cdc42 as a hub^40,41^. Thus, loss of *Sarm1* in stroke-injured cortical neurons results in selective transcriptional expression of genes with known, pro-growth functions including axonal transport, growth and repair.

**Table 1.**
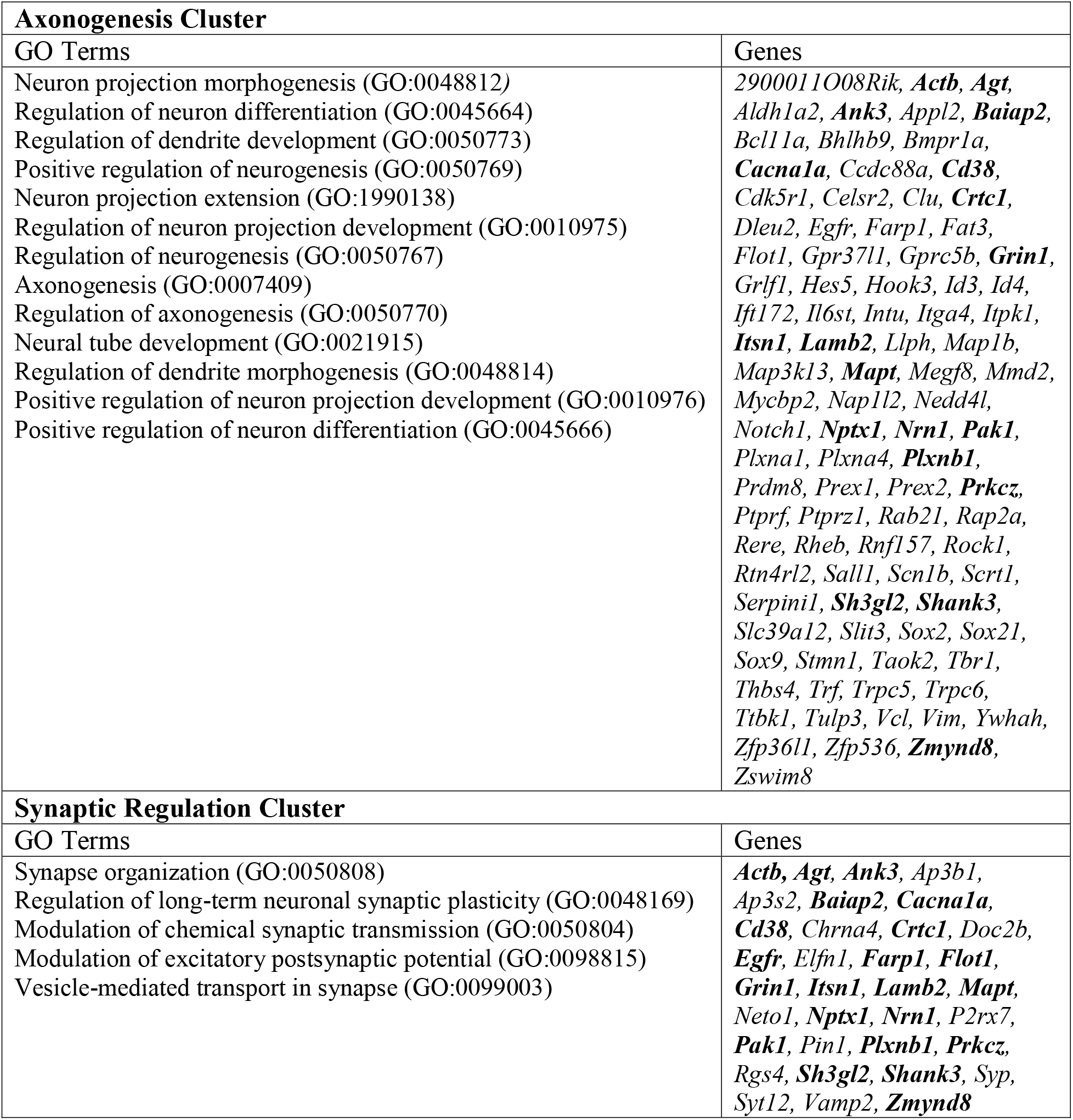
Differentially regulated genes in axonogenesis and synaptogenesis clusters. Bold indicates genes shared in both clusters. All GO terms were significant at *p*<7.28x10^-4^. Differential gene expression threshold defined at FDR<0.05.

### Functional genomics identifies small molecular compounds that recapitulate the molecular signature induced by loss of *Sarm1* in stroke-injured neurons and promote neurite growth

We further reasoned that this novel regenerative program manifested by the loss of *Sarm1* in cortical neurons could be exploited to identify novel small molecule therapeutic candidates to replicate the pro-growth transcriptional program and potentially promote axonal sprouting or remodeling after ischemic stroke. To achieve this, we conducted a functional genomics screen using the top 150 differentially regulated genes between *Sarm1-/-* vs WT stroke-injured cortical neurons (Fig. 5A). Using the CLUE-IO bioinformatics database^42^, we identified 2428 perturbagens capable of mimicking or inhibiting the transcriptional signature of *Sarm1-/-* stroke-injured cortical neurons (see also Table S4). Employing a structured filtering strategy (Fig. 5B), we refined this compound library down to 18 molecules with known or predicted blood-brain barrier permeability. Several different pharmacologic drug classes were included among the identified compound library including adrenergic and androgen receptor antagonists, neurotransmitter (DA, 5-HT) agonists and antagonists, as well as enzymatic inhibitors and DNA modulators (Fig. 5B).

**Figure 5.**
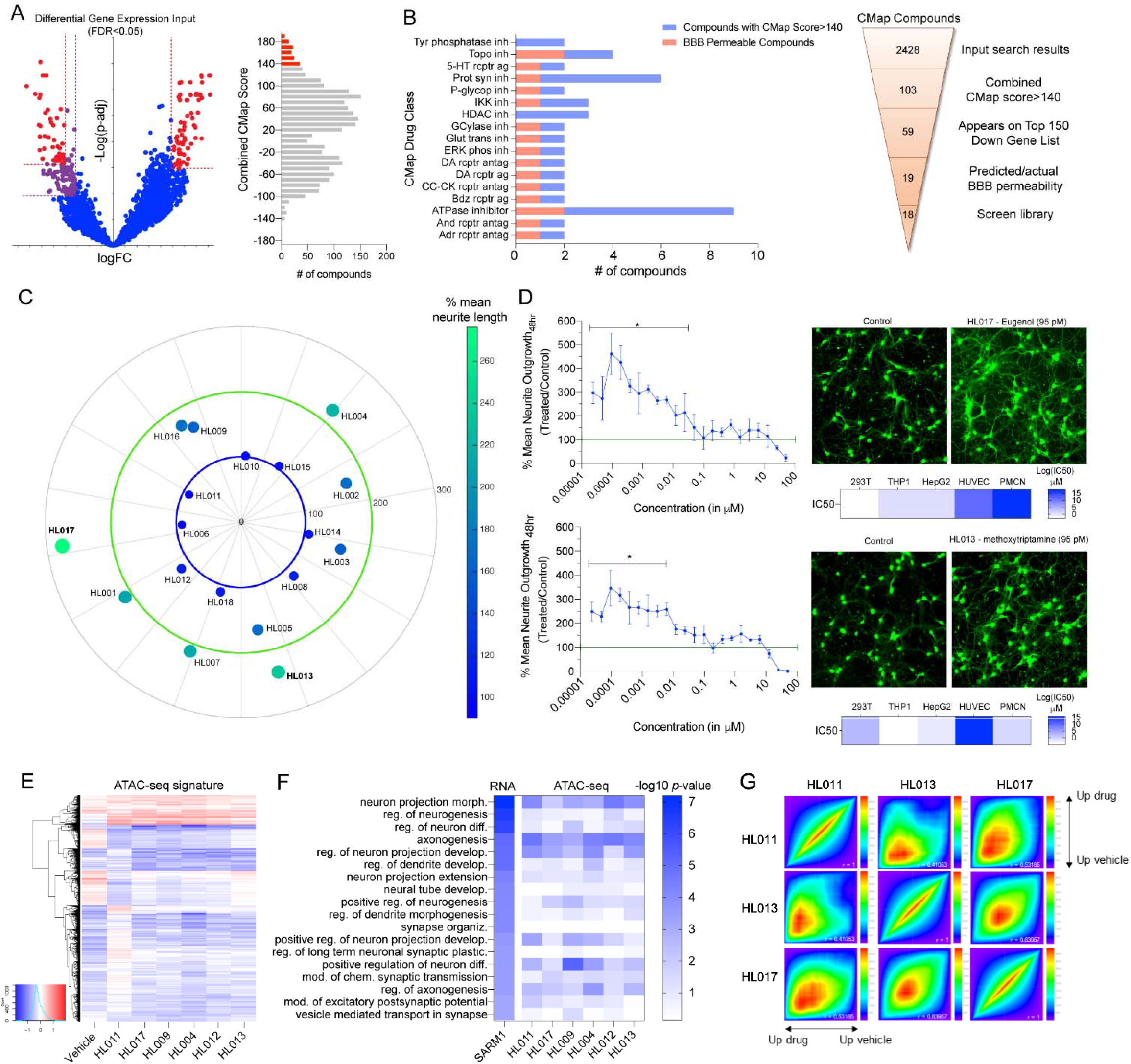
Functional genomics screen for pharmacologic targets to replicate axonogenesis and synaptogenesis programs identified in stroke-injured *Sarm1-/-* cortical neurons. Volcano plot of top differentially regulated genes in *Sarm1-/-* stroke-injured cortical neurons (red = top 150; purple = top 150 down-regulated) used in CLUE-IO query to identify compounds with combined CMap threshold score ≥ 140 (red) (A). Bar plot showing number of compounds with CMap score ≥ 140 by pharmacologic class (blue) and blood-brain barrier permeability (red) (right) with schematic of filtering strategy for compound library generation (right) (B). Polar plot of neurite outgrowth (% mean neurite outgrowth at sub-nanomolar doses) by each compound tested vs. control cortical neurons (three replicates/drug/dose) (C). Dose effect of neurite outgrowth for two hit compounds (HL017 – Eugenol; HL013 – 5-methoxytryptamine) (*p*<0.0001 by two-way ANOVA; *adjusted *p*<0.01 for individual dose comparisons vs. control, *n*=4 replicate cultures/dose) with representative Calcein-AM imaging. Horizontal heatmaps of logIC_50_ (μM) across five cell lines (D). Neurite outgrowth of cortical neurons following rotenone-induced ischemic injury and cultured with either one of the two hit compounds (HL013, HL017) vs vehicle is further shown in Figure S5. Heatmap of ATAC-seq signature from drug-treated E18 mouse cortical neurons (FDR<0.05) (E). Heatmap of up-regulated GO terms vs. vehicle identified by ATAC-seq in drug-treated neurons plotted vs. GO terms from *Sarm1*-/- axonogenesis and synaptogenesis clusters (F). Coordinated epigenetic signature of neurons treated with the compounds identified from the functional genomic screen is further shown in Figure S6. RRHO heatmaps of ATAC-seq signatures of drug-treated E18 mouse cortical neurons comparing growth-promoting (HL013 and HL017) and non-growth promoting (HL011) compounds (G). Shared transcription factor motif analysis from neurons treated with the pro-neurite regrowth compounds as identified from the functional genomic screen is further shown in Figure S7. Error bars = SEM.

Next, using a high-throughput drug screen design employing murine E18 cultured cortical neurons, we analyzed the extent of neurite outgrowth promoted by this refined library of 17 compounds (excluding one Schedule II FDA-regulated drug). Beginning at DIV7, we exposed wild-type cortical neurons to library compounds for 48 hrs across a 20-step dilutional curve. Immediately after drug exposure, we measured mean neurite outgrowth at doses below 100 nM. We identified 5 of 18 compounds that resulted in greater than 2X of neurite outgrowth in vitro (hypergeometric *p* = 0.024; Fig. 5C). For the majority of hit compounds, drug effects on neurite outgrowth were prominent at the low end of the dilutional curve. Moreover, among the hit compounds, HL017, the androgen receptor antagonist eugenol, demonstrated robust neurite outgrowth in the sub-nanomolar range while having a fairly high IC50 for toxicity in primary cortical neurons (*p*<0.0001 by two-way ANOVA, F_(21, 682)_=19.32, logIC_50(PMCN)_=17.72 μM, Fig. 4D). Similarly, HL013, the 5-HT receptor agonist 5-methoxytriptamine, also drives neurite outgrowth at very low doses while having a favorable toxicity profile in primary cortical neurons (*p*<0.0001 by two-way ANOVA, F_(21, 682)_=13.47, logIC_50(PMCN)_=-1.37 μM, Fig. 5D). To determine if these hit compounds could also promote neurite re-growth after ischemic injury, we modeled chemical ischemia *in vitro* by treating the primary cortical neurons with rotenone for 3 hours, followed immediately by treatment with either HL013 or HL017 compound. In this chemical ischemia model, both HL013 and HL017 compounds significantly increased neurite re-growth after rotenone-induced ischemic injury as compared to vehicle treated neurons (*p*=0.0005 by ordinary two-way ANOVA, F_(2, 8)_=22.74; see also Fig S5). Thus, using functional genomics we identified compounds that independently mimic the pro-growth molecular profile of stroke-injured *Sarm1-/-* neurons, and moreover are sufficient to promote endogenous neurite outgrowth as well as neurite re-growth after ischemic injury *in vitro*.

### Molecular candidates recapitulate the pro-growth molecular program in *Sarm1-/-* cortical neurons by modulating chromatin accessibility

Because the known mechanism of action of all five hit compounds was substantially different from each other and their maximal effect on neurite outgrowth was most prominent at low dose, we postulated that these compounds may be working to drive axonal outgrowth via a novel mechanism. Given that these compounds were all identified using the CLUE-IO resource employing a design to mimic a pro-regenerative transcriptional profile, the most reasonable shared mechanism among these compounds was via epigenetic regulation of the neuronal genome. To establish this novel mechanism, we exposed E18 wild-type cortical neurons to the hit compounds with the maximally effective dose driving neurite outgrowth for 48 hours, and then performed ATAC-seq to assess the degree of genome wide chromatin accessibility and the expression changes of accessible genes.

Compared to vehicle controls and a screened compound that did not drive neurite outgrowth in vitro (HL011), all five hit compounds generate a similar epigenetic signature of differentially accessible genes (DAGs) in cortical neurons (Fig. 5E; and see also Fig. S6 and Table S5). Notably, the significantly enriched GO terms that comprise the axonogenesis and synaptic regulation clusters as identified in *Sarm1-/-* stroke-injured cortical neurons by RNA-seq are also conserved in the up-regulated GO terms derived from DAGs by ATAC-seq across all five hit compounds that promote neurite outgrowth *in vitro* as compared to vehicle-treated cells (Fig. 5F).

To statistically define similarities in genome-wide chromatin accessibility driven by the hit compounds, we further compared the gene accessibility profiles between non-growth promoting hit compounds (eg: HL011) and growth promoting hit compounds (eg: HL013, HL017) using the rank-rank hypergeometric overlap (RRHO) algorithm (Fig. 5G). RRHO heatmaps confirm greater similarity in gene accessibility signatures among the hit compounds that show growth-promoting properties, with compounds HL013 and HL017 demonstrating the highest level of genome-wide similarity (*r*=0.64, *p*<0.0001). Conversely, DAG comparisons amongst non-growth promoting compounds including HL011 (*r*=0.41, *p*<0.0001) and HL013 (*r*=0.53, *p*<0.0001) were still significant but not as strong, indicating coordinated and specific epigenetic signatures driven primarily by the growth-promoting hit compounds. Motif analysis further demonstrates 8 up-regulated and 2 down-regulated transcription factor recognition motifs that were enriched and conserved across all the growth promoting hit compounds compared to vehicle and non-growth promoting compounds (HL011) (see also Fig. S7).

Notably, the corresponding transcription factors (*RBPJ, Ascl, E2A*) containing these conserved motifs all have recognized roles in neuronal- and axonal-genesis^43,44,45,46^, thus further reflecting an epigenetic control of a pro-growth molecular program by Sarm1 in cortical neurons.

## DISCUSSION

Using a well-characterized *in vivo* murine model of focal white matter stroke, we show that loss of Sarm1 significantly promotes long distance and long-acting (up to 28 days--the latest time point after stroke that we evaluated) structural preservation of the axons after focal ischemic stroke to the white matter. Our data definitively demonstrate that Sarm1 is required for axonal degeneration from ischemic injury, and that axonal degradation following ischemia, similar to multiple types of traumatic and chemotoxic mechanisms of CNS injuries, converge on a Sarm1-dependent mechanism. Thus, interventions aimed at inhibiting or abrogating Sarm1 activity present a viable and effective strategy to mitigate axonal degeneration after ischemic stroke.

We further demonstrate that the absence of Sarm1 also results in a significantly increased number of surviving neurons distant from the infarct in two different ischemic stroke models. This includes increased survival of cortical neurons following a subcortical white matter stroke, and increased survival of subcortical thalamic neurons labelled with an AAV-DJ-hSyn-EGFP viral reporter after a cortical stroke by permanent occlusion of the distal middle cerebral artery. Given that the AAV-DJ virus used to label the thalamic neurons are not known to be capable of transsynaptic transmission, the labelled subcortical neurons are most likely thalamocortical neurons with axon terminals in the cortex retrogradely labeled at the site of ischemia through a mechanism that is well described for AAV vectors^63,64,65^. In both cases, there is a significantly greater number of stroke-injured but surviving neurons in *Sarm1-/-* mice compared to WT mice, and this neuroprotection occurs even in regions remote from the site of stroke lesion (but along the expected paths of neurite projections and/or synaptic targets of the stroke-injured neurons), with increased number of surviving neurons distant from the infarct in two different ischemic stroke models. Importantly, we further show that absence of Sarm1 results in significant improvement in motor and cognitive deficits following white matter stroke, with continued improvement in both motor and cognitive functions up to 30 days post-stroke. Thus, our results indicate that loss of Sarm1 also mitigates secondary degeneration of neurons that are distant from but reasonably assumed to be anatomically connected to the infarct lesion^47, 48^ after stroke, and moreover this neuroprotection leads to significant clinical recovery and amelioration of functional outcomes after ischemic stroke.

Our data further suggests that a parsimonious mechanism by which loss of Sarm1 mitigates such secondary axonal and neuronal degeneration, and promote stroke recovery may be by preventing the activation or transport of injury signal(s) from the stroke-injured synapse or axon to the soma that then trigger eventual degeneration of the cell body. Indeed, our finding that loss of Sarm1 promotes the survival of neurons remote from but connected to the stroke lesion further supports a “network effect” mechanism in causing secondary neurodegeneration after stroke^49^, and that *Sarm1-/-* mediated axonal preservation may be preventing injury signals from triggering or propagating further degeneration of neurons distant from the site of stroke injury. However, we are unable to resolve more precisely whether the surviving *Sarm1-/-* neurons that are distant from the infarct site are mostly second order, post-synaptic neurons distal from the neurons with their cell bodies originating at the stroke lesion (and their axons protected from anterograde axonal degeneration), or are neurons with cell bodies at the distal site and send neurite projections to or synapses at the stroke lesion (and their axons protected from retrograde axonal dieback). More detailed dual labeling and/or trans-synaptic labeling (ie, rabies virus based labeling) strategies to delineate the underlying circuitry connecting the stroke lesion and the distally protected *Sarm1-/-* neuronal cell bodies may further reveal the directionality and nature of secondary neurodegeneration.

Notably, several studies have already identified candidate “axonal injury signals” such as c-Jun N-terminal kinases (JNK), a member of the MAPK family, to be retrogradely transported from the axon to the soma and modulate the expression of known injury response molecules following traumatic peripheral nerve as well as optic nerve injuries^50^. Moreover, downstream MAPK kinases including MAPK3, DLK and LZK are implicated in activating JNK1–3 and their canonical target JUN, which leads to transcriptional changes that ultimately culminate in BAX activation and neuronal apoptosis^50, 51^. Curiously, loss of Sarm1 delayed axonal degeneration, but did not prevent DLK/JNK pathway activation or delay RGC cell death in traumatic optic nerve crush^52^. In contrast, we observed in stroke-injured FACS-captured cortical neurons limited transcriptional activation of Mapk kinases in *Sarm1-/-* mice compared to wild-type mice. Thus, while Sarm1 mediates a common mechanism of axon degeneration across multiple modes of injury, there may be heterogeneity in the injury signals involved in triggering or causing subsequent neuronal degeneration. This may be further resolved with additional studies ascertaining whether the JNK/MAPK pathway are specifically activated after stroke in our *in vivo* models, and whether a causal relationship exists between Sarm1 activity and activation of retrograde axonal injury signals following ischemic injury.

To further determine the downstream mechanism of *Sarm1-/-* mediated neuronal protection, we compared differential gene expressions of stroke-injured WT and *Sarm1-/-* neurons and unexpectedly identify a restricted subset of DEGs in stroked-injured *Sarm1-/-* neurons that cluster to pro-growth functional groups. Specifically, we identified unique molecular signatures in stroke-injured neurons deficient in *Sarm1* that favor a pro-growth pattern of expression and support post-stroke neural survival and repair. The restricted number of DEGs and the selectivity of the transcriptional response further indicate the internal validity of our transcriptomic analyses, as a generalized, non-specific consequence of Sarm1 loss (e.g., a global injury or stress response) would be expected to produce broad, largely non-specific changes in gene expression. Instead, we observe that only a restricted subset of genes enriched for functions in axonal transport, growth, and repair changes in a Sarm1-genotype-dependent manner.

Notably, our transcriptomic analyses between *Sarm1-/-* and WT stroke-injured cortical neurons did not show enrichment or alteration of DEGs that are normally known to participate in previously defined Sarm1 signaling pathway^35,36^. This indicates that the transcriptional responses of cortical neurons to ischemic injury in the absence of *Sarm1* involve specific expression changes of novel pathways distinct from known Sarm1 targets.

As recovery and neural repair after stroke is ultimately mediated by the development of new axonal connections between disconnected brain regions, our findings suggest that in addition to its local activity in regulating axon degeneration after stroke, Sarm1 and/or its downstream target(s) normally inhibit a robust pro-growth axonogenesis program in cortical neurons after stroke. In fact, several pathways contributing to axon degeneration overlap in part with those that drive axon regrowth^53, 54^, suggesting a convergence of mechanisms that regulate axonal degeneration and attempted axonal repair in the CNS after injury^55^. In our study, the expression of pro-growth genes promoted by the loss of the degenerative molecule Sarm1 further supports such an overlap between axonal degeneration and inhibition of reparative programs for neural modeling after stroke, and raises the prospect that axonal degeneration and axonal remodeling after stroke injury are both regulated by Sarm1 activity. However, whether this pro-reparative program induced by loss of Sarm1 is directly mediated by Sarm1 activity itself or indirectly by downstream targets of Sarm1 remains unclear.

Mechanistically, we further showed that this axonal pro-growth program in cortical neurons that is normally inhibited by Sarm1 activity is regulated at the level of the epigenome. Previous studies have shown that regulation of NAD+ availability modulates gene expression changes via indirect actions on the chromatin or the transcriptional machinery. Notably, several NAD -dependent enzymes — most notably the sirtuins, including Sirt1, as well as PARP, an NAD+ dependent ADP-ribosylase—act as downstream molecular effectors of NAD+ and couple cellular NAD status to gene expression via deacetylation or ADP-ribosylation of chromatin histones and transcription factors, respectively^67,68^. Thus, Sarm1 may modulate this pro-growth transcriptional program epigenetically through its NADase activity and control of NAD availability which then leads to differential activities of NAD+ dependent, chromatin-facing proteins. This model is consistent with prior work showing that (i) NAD availability acting through Sirt1 and NAD pathway enzymes directly regulates chromatin state and target gene transcription (ii) manipulating nuclear NAD biosynthesis activity is sufficient to alter neuronal/axonal survival and gene expression programs, and (iii) Sarm1 has established roles not only in axon degeneration but also in regulating gene transcription and axon regeneration through NAD -linked and NAD -independent signaling arms^66,67,68^. Further experiments to address this hypothesis, such as manipulating Sirt1 or PARP activity independently of Sarm1 genotype, will be needed to establish causality, and a potential future direction based on our data.

Lastly, we identified molecular compounds from our functional genomics screen that recapitulate a similar pro-growth genetic program favoring axonogenesis as that seen in *Sarm1-/-* stroke injured neurons, and we showed that these compounds independently promote *de novo* axonal outgrowth as well as regrowth following ischemic stressor conditions *in vitro*. Although the pro-growth molecular compounds identified from the screen individually target different mechanisms, the presence of conserved motifs that are similar between DAGs in ATAC-seq and bulk RNAseq between WT and *Sarm1-/-* neurons indicates a highly conserved, de novo epigenetic pro-growth program in cortical neurons that is druggable by modulating differential accessibility to chromatin at defined loci. Thus, our study indicates that pharmacologic interventions to modulate Sarm1 activity itself, or to phenocopy the same pro-growth expression profile as *Sarm1-/-* in cortical neurons after stroke may concurrently attenuate axon degeneration and promote the reparative capacity of cortical neurons after ischemia, and altogether points to a potential new approach to enhance neural repair after stroke by targeting Sarm1.

In summary, we demonstrate that loss of Sarm1 mitigates anterograde axonal degeneration and promotes neuronal survival, and this leads to significant ameliorations of motor as well as cognitive functions after ischemic stroke. We further show that loss of Sarm1 facilitates a reparative program in cortical neurons that is encoded at the epigenetic level. Our finding that this epigenetic signature can be recapitulated with multiple low dose pharmacologic agents supports the concept that a regenerative capacity for axonal outgrowth in cortical neurons after stroke is programmed into the genome but actively restricted by pro-degenerative regulators such as Sarm1. Well-timed inhibition of Sarm1 to relieve this regulation, or direct activation of this downstream, druggable epigenetic program may thus represent a novel strategy to mitigate axonal degeneration, limit secondary neurodegeneration and promote axon repair after stroke.

## MATERIALS AND METHODS

### Animals

All animal studies presented here were approved by the UCLA and Stanford University Animal Research Committees, both accredited by the AAALAC. Mice were housed under respective institutional regulation with a 12-h dark-light cycle. All mice used in the study were male between the ages of 3-6 months. Strain matched wild-type C57BL/6 mice (Jackson Labs, Strain #000667) and *sarm1-/-* mice (Jackson Labs, Strain #018069) backcrossed and bred in C57BL/6 background x 3 generations were used for all experiments unless otherwise stated.

### White matter ischemic stroke

Subcortical white matter ischemic injury was induced as previously described^6,28^ using focal injections into subcortical white matter x 3 of the iNOS inhibitor L-N^5^-(1-Iminoethyl) ornithine, Dihydrochloride (LNiO; 27 mg/mL, Millipore) added at 1:1 ratio with 20% fluororuby (Fluorochrome LLC) dissolved in saline to label stroke-injured axons and connected cortical neurons. Animals were sacrificed at 7 and 28 days after stroke, and freshly dissected and transcardially perfused with 4% PFA and prepared for tissue sections as previously described^6^.

### AAV labeling of cortical neurons

Stock adeno-associated virus encoding the EGFP reporter under the control of hSyn neuronal promoter (AAV-DJ-hsyn-EGFP) was obtained from the Stanford Gene Vector and Viral Core (GVVC-AAV-127). The AAV-DJ-hsyn-EGFP construct was then stereotactically injected into the cortex of WT and *sarm1*-/- mice. Specifically, an 1 μL microsyringe (Hamilton, Cinnaminson, NJ) was used inject the virus (2 μL;1E+12v.g./mL) into the cortex at anticipated location of the infarct from permanent distal MCA occlusion (see below) using the following stereotactic coordinates: -0.26mm posterior and +1.3mm lateral with respect to the bregma, and a depth of -2.17mm from the surface of the cortex. To allow sufficient gene expression, all injections were completed 2 weeks before cortical MCAO stroke surgery as described below.

### Distal middle cerebral artery occlusion (dMCAO) stroke

Ischemic stroke to the cortex via permanent dMCAO was performed as described previously^29,30^. In brief, mice were anesthetized by isoflurane inhalation (2% isoflurane in 100% oxygen) and an incision was made to expose the right temporalis muscle. A pocket was created in the temporalis muscle and the right middle cerebral artery (MCA) was identified through the skull underneath. A microdrill was used to penetrate the skull and expose the underlying MCA branch. The meninges were then cut, and the distal MCA vessel was cauterized using a high-temperature Bovie (Ambler Surgical, PA). After complete occlusion of the underlying MCA was visually confirmed, the wound was closed using surgical adhesives. Throughout surgery, the animal body temperature was maintained at 37°C using a feedback-controlled heating blanket.

### Grid-walking task

Motor deficits were assessed using the Foot Fault Scan system (CleverSys). The apparatus consists of a 12-mm square wire mesh (32 × 20 × 50 cm, L × W × H) elevated above a laser beam network and video camera to detect stepping errors. Mice were placed individually on the grid and allowed to explore freely for 5 min. Foot faults (steps failing to provide support, with the paw passing through the grid) and non-foot fault steps were recorded for each limb. Percent foot faults were calculated as:

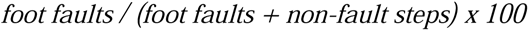

Foot faults were normalized to total steps to control for differences in locomotor activity across animals and trials.

### Spontaneous forelimb task (cylinder task)

Forelimb asymmetry was evaluated as previously described^62^. Mice were placed in a transparent Plexiglas cylinder and recorded for 5 min during spontaneous rearing and wall exploration. Videos were analyzed in slow motion (1/5th real time speed) by a blinded observer. During vertical exploratory movements in which both forelimbs were clearly visible, contacts made with the left (contralateral) forelimb, right (ipsilateral) forelimb, or both forelimbs were counted. A spontaneous forelimb asymmetry index was calculated as:

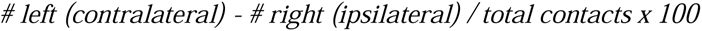

### Fear conditioning task

Associative learning and memory were assessed using a standard cued fear conditioning paradigm^31^. During training, mice received three pairings of tone/light cues with a 2-s, 0.75-mA foot shock. For retrieval 24 h later, mice were re-exposed to the same cues without shock. Freezing behavior was automatically quantified (Freeze Scan, CleverSys) and expressed as percent time freezing. Memory performance was calculated as the change in freezing between retrieval and training sessions (% retrieval − % training).

### Immunofluorescence and confocal imaging

Fluororuby-labeled WT and *Sarm1-/-* stroke-injured brains were harvested, perfused and cryosectioned at 40 μm in a -22C cryostat and then stored in cryoprotectant at -20C. For staining, tissue sections containing stroke were removed from the cryoprotectant and washed in PBS and incubated for 30 min in 10 mM sodium citrate buffer. After cooling and washing, sections were blocked in PBTDS and the tissue was incubated overnight in anti-NF-H (Sigma, 1:500) to immunostain for neurites, anti-NeuN (Abcam, 1:500) for neuronal cell bodies, and anti-CD68 (Abcam, 1:1000) for macrophages (to delineate the stroke core and peri-infarct/penumbral regions after dMCAO). Corresponding secondary antibodies were added (1:500) including Donkey anti-Rabbit 488, Alexa Fluor 555-conjugated goat anti-rabbit IgG (1:500; Abcam), Donkey anti-Mouse 647 (Jackson ImmunoResearch) and counterstained with DAPI. The tissue was mounted onto glass slides and dehydrated in ethanol and xylenes and covered with DPX and a coverslip. Imaging was conducted on a Nikon C2 confocal microscope. Three 60X images were taken on each tissue section in the region of interest containing stroke-injured and non-stroke injured axons or cortical neurons.

Axonal morphology and integrity were evaluated using ImageJ/Fiji software (NIH). The axons were measured by manual tracing beginning at infarct core and moving medially towards midline of corpus callosum in 50um increments. Fluorescence intensity within the corpus callosum as manually traced in each ROI was then quantified by ImageJ/Fiji software.

For all confocal imaging of thalamic neurons, anatomical landmarks were used to identify the thalamus (59). This included first the selection of stroked tissue sections (marked by +CD68 staining) with the hippocampus in view. This is then followed by identification of the region of intersection between the hemispheric midline and the corpus callosum. Thalamic region was then characterized in all sections as 2 fields of view (FOV) lateral to the midline and 3 FOV inferior to the corpus callosum on a 40X mag view.

### u-DISCO tissue clearing and neuronal density measurement

At 7 days after white matter stroke, brain tissues of WT and *Sarm1-/-* animals (*n* = 4 in each cohort) were transcardially perfused with 4% PFA and post-fixed overnight at 4C. Tissue slabs 3 mm in thickness and spanning the region of stroke were generated that included left and right cortical regions. Tissues were cleared using uDISCO as previously described^33^. Briefly, tissues were optically cleared by serial incubation in increasing concentrations of tert-butanol (Acros Organics) followed by immersion in benzyl alcohol (Sigma-Aldrich)/benzyl benzoate (Sigma-Aldrich)/diphenyl ether (Alfa Aesar) (BABB-D) solution until transparent. The tissues were then immediately imaged on a Leica SP5 laser confocal microscope. Cell density was analyzed and quantified using Imaris software. A 3D grid was applied to images in the contralateral cortical hemisphere and the density of FR+ neurons within each field of view was measured. Computer-generated labeling of each cell body based on cell body size, diameter and circularity was performed to aid in the identification and quantification of the neuronal cells.

### Stroke-injured neuron isolation by MACS-FACS

At 7 days after focal white matter stroke, regions of cortex ipsilateral to and overlying the subcortical white matter stroke lesions were dissected and mechanically digested following a single-cell suspension protocol using a Neurocult dissociation kit (STEMCELL) from 4 animals per genotype. For magnetic bead cell sorting (MACS), neuronal enrichment kit microbeads and CD11b microbeads (Miltenyi Biotec) were added to the suspension before applying to a MACS column to negatively select non-neuronal cells. After collecting all neuronal cells, cells were further labeled with anti-NCAM antibody (Abcam) followed by anti-rabbit Alexa 488 secondary antibody to label cortical neurons and sorted by FACS for fluororuby and NCAM+/Alexa488+ neurons at the UCLA Flow Cell Cytometry Core. Total RNA was collected from sorted cells using conventional RNA preparation kit (RNeasy, Qiagen).

### Gene expression profiling

Isolated RNA from MACS-FACS isolated cortical neurons was normalized by FACS cell counts. Harvested RNA samples at 0.5ng/ul concentration were validated and quality-controlled using TapeStation 2200 bioanalyzer (Agilent Technology). cDNA library creation performed by the UCLA Neuroscience Genomic Core (UNGC) using the NuGEN Ovation Ultralow system with KAPA HyperPrep and sequencing on the Illumina HiSeq 4000 platform. The quality of sequencing was assessed using FastQC, followed by read alignment with STAR aligner. Differential gene expression was performed using DESeq2. After read-count normalization, differentially expressed genes (DEGs) with FDR<0.05 were then compared. Gene ontology analysis was performed by comparing to the whole mouse genome using GOrilla^56^. Resulting gene ontology terms (p<0.05) were used as input to REVIGO^57^. Semantic clustering of gene ontology terms derived from REVIGO were plotted via MatLab to create a gene ontology map. The list of all DEG and GO terms used are provided in Table S3.

### Protein-protein interaction network analysis

Protein-protein network interactions were evaluated using the STRING Db resource from 2019. Conserved genes in both the axonogenesis and synaptic regulation GO terms were used as input criteria against the whole genome with the following search parameters: evidence for network edges; all sources; high confidence 0.7; no interactors.

### Functional genomics screen

The top 150 differentially expressed genes by log-fold change and the top 150 down-regulated genes by log-fold change were used as individual gene expression (L1000) query inputs for CLUE-IO in two separate trials using the latest Touchstone database (2018). Queries were compared to reference perturbagens in cell lines including human melanoma (A375), human non-small cell carcinoma (A549), human lung adenocarcinoma (HCC515), human liver cancer (HEPG2), human colorectal adenocarcinoma (HT29), human breast cancer (MCF7), human prostate cancer (PC3), immortalized human kidney (HA1E), and human adherent, epithelial prostate cancer (VCAP) cell lines (https://clue.io/). Compound lists were generated for each trial producing a total of 2428 compounds. CMap scores were summated, and compounds with a summated enrichment score ≥140 were curated and refined further based on actual or predicted blood brain barrier permeability by literature search, resulting in the identification of 18 viable candidates labeled with the identifiers HL001 through HL018.

### Cell culture

Primary mouse cortical neurons (Thermo Fisher Scientific, Catalog #A15586) were plated at a density of 5000 cells/well and cultured in Neurobasal media (Gibco, Ref #21103-049) supplemented with B27 (Gibco, Ref #17504044) and Glutamax (Gibco, Ref #35050-061) in black 384 well flat bottom microClear cell culture microplates (Greiner Bio One, Cat #781091). Prior to plating, the culture plates were coated with 50µg/mL of Poly-D-Lysine solution (Cat #A3890401). Plates were incubated overnight in 37°C, 5% CO_2_ humidified incubator. Poly-D-Lysine solution was aspirated prior to washing wells 3x with sterile dH_2_O and aspirating wells completely to dry. Neurons were cultured for seven days with half-media exchanges every 2-3 days. Chemical ischemia was induced by exposing the cultured neurons to 3 hrs of rotenone treatment (25μM in 0.5% DMSO) as previously described (Sur et al, 2018). For drug toxicity measurement, four additional cells lines (293T, THP1, HepG2, and HUVEC) were cultured in normal growth conditions for 48-72 hours in CELLSTAR uClear 384 well plates at seeding density of 2000 cells/well in respective growth media.

### Pharmacologic screen

The final compound library set was 17 compounds due to elimination of Alprazolam as a DEA Schedule 4 drug. All compounds were diluted in 0.5% DMSO (Sigma, Cat #276855) at 10 μM. Diluted compounds were pinned to 384 well plates with 25µL of cell media per well with pin size of 250nL using BioMek FX (Beckman Coulter, CA). DIV7 cultured neurons plated in 384 well plates were exposed to compounds across a 20-step serial dilution (24pM-50µM) for 48 hours in a 37°C, 5% CO_2_ humidified incubator without media exchange. Cellular toxicity was established using propidium iodide (Invitrogen, Cat #P3566) assay. To determine the effects of compound exposure on neurite outgrowth, cells were exposed to individual compound for 48 hrs, washed three times, and loaded with 1µM Calcein-AM (Sigma, Cat #17783-1MG) and Hoechst (Invitrogen, Ref #H3570) before a single well FOV was captured on an ImageXpress Confocal (Molecular Devices, CA) using a 10x objective. All experiments were conducted in triplicates. Total neurite outgrowth was measured via a MetaXpress (Molecular Devices, CA) neurite outgrowth analysis algorithm. Neurite outgrowth was set to detect cell bodies with approximate width of at least 20µm and outgrowth with maximal width of 5µm and length of at least 100µm with intensity of 1000 grey scales over background. Mean neurite outgrowth per drug exposure was determined by normalizing to control (*n*=30).

### Assay for transposase-accessible chromatin with high-throughput sequencing

Primary mouse cortical neurons (Thermo Fisher Scientific, Catalog #A15586) were thawed and seeded in a 24 well TC treated plate coated with Poly-D-Lysine and Laminin. Neurons were grown in 24 wells at seeding density of ∼475k cells/well. Cells were fed every 2-3 days with half media changes. On day 6 of cell culture, neurons showed axonal processes and were exposed to 8 treatment conditions: PMCN, Vehicle, HL004, HL009, HL011, HL012, HL013, and HL017. The neurons were incubated in each condition for 48 hours. Each condition was run in triplicates. After drug incubation, the neurons were then lysed on the plate using the ATAC-RSB-Lysis buffer and collected with a cell scraper to count ∼50k nuclei from each well per condition on a hemocytometer with trypan blue staining. The isolated nuclei were prepared for transposition by incubating with the transposase enzyme Nextera Tn5 Transposase (Illumina, REF #20034210) and then purified with the Qiagen MinElute PCR Purification kit (Qiagen, REF #28004). Transposed DNA was eluted in 10µL and subjected to PCR amplification and library generation in the UCLA Neurogenetics and Genomics Core Facility. Resulting DNA sequences were aligned to the mouse genome (mm10) using Burrows-Wheeler Aligner mem with default parameters. Duplicates were removed with Picard software. Reads were pre-shifted by 75 bp prior to peak calling using MACS2. Gene ontology analysis was further performed and refined using Homer and conserved motifs across the vehicle control, negative control, and 5 drug conditions that were thresholded to remove motifs enriched in vehicle and HL011 (no growth control). The remaining motifs were sorted by significance to identify top hits related to drug treatment.

### Rank-rank hypergeometric overlap (RRHO) analysis

Gene accessibility signatures derived from ATAC-seq were compared using the RRHO algorithm (58). Each signature was processed as a ranked list using different expressions for two classes of compounds. Signature inputs were pre ranked gene lists with peak ID, gene name, rank 1, rank 2, metric 1, and metric 2. Gene lists were pre ranked greatest to least logFC according to the first compound in the comparison serving as rank 1 and metric 1. For metric 1 and 2, the logFC values of compound comparisons were used with FDR 0.05. Step size of 100 was used to bin the ranked items to improve the run time of calculating the hypergeometric distribution. R values were calculated as a correlation between the comparisons of two compounds.

### Statistical analysis

Imaging analysis and data quantification were performed using ImageJ (NIH) and Prism v7.0 software (GraphPad). Error bars shown in all graphs are standard error of the mean (SEM). Gene expression and gene accessibility values were normalized and compared using a false-discovery rate adjusted *p*-value assuming significance at FDR < 0.05. A paired two-tailed t-test was used to compare the intensity of fluororuby-labelled WT and *Sarm1-/-* axons, and to compare the mean number of NeuN+ neurons in the thalamus between WT and *Sarm1-/-* animals. Unpaired two-tailed t-test was used to compare neuron counts in 3D uDISCO analysis. One-way ANOVAs were used to verify neuronal and layer-specific enrichment. Neurite outgrowth assay data were compared using a two-way ANOVA for drug effect and concentration. All judgements of statistical significance were performed using post-hoc adjustments for multiple comparisons with a starting assumption of α=0.05.

## RESOURCE AVAILABILITY **/** DATA SHARING STATEMENT

### Lead Contact Statement

Requests for further information and resources should be directed to and will be fulfilled by the corresponding author.

### Materials Availability

All unique/stable reagents generated in this study are available from the lead contact with a completed materials transfer agreement.

## Data and Code Availability

1) The gene expression and ATAC-seq datasets generated during and/or analyzed during the current study are available in the GEO repository (ascension number GSE280862; https://www.ncbi.nlm.nih.gov/geo/query/acc.cgi?acc=GSE280862) and are publicly available as of the date of publication.

2) Additional imaging data is available on the following Open Science Framework weblink: https://osf.io/7gqvt/?view_only=f643d69a4f224fc7a356c9d71a2a31f3

### Funding

This work was generously supported by NINDS R25NS065723 (JTW), K08 NS083740 (JDH), UC Drug Discovery Consortium (JDH, RD).

