## Supplemental Information for "Loss of Sarm1 Mitigates Axonal Degeneration and Promotes Neuronal Repair After Ischemic Stroke"

**Supporting Information Appendix**

Figures: 7

Tables: 5

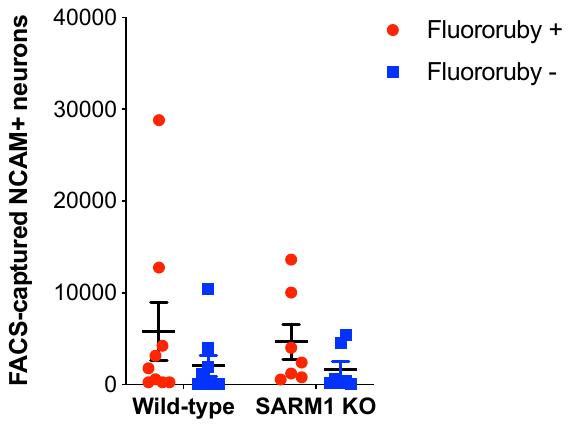

**Figure S1. MACS-FACS isolation of stroke-injured cortical neurons.** Seven days following subcortical ischemic stroke, sensorimotor cortex overlying the stroke lesion was dissected and subjected to MACS to remove non-neuronal cells. Resulting cell suspension was labeled with anti-NCAM-488 and NCAM+/fluororuby+ cells (red) were selected by FACS in both wild-type and SARM1 knockout mice (*n*=9 per group). Equal numbers of NCAM+/FR+ cells were sorted in both groups (*p*=0.87, F_(1,28)_=0.03 by two-way ANOVA; adjusted *p*=0.98 for genotype effect).

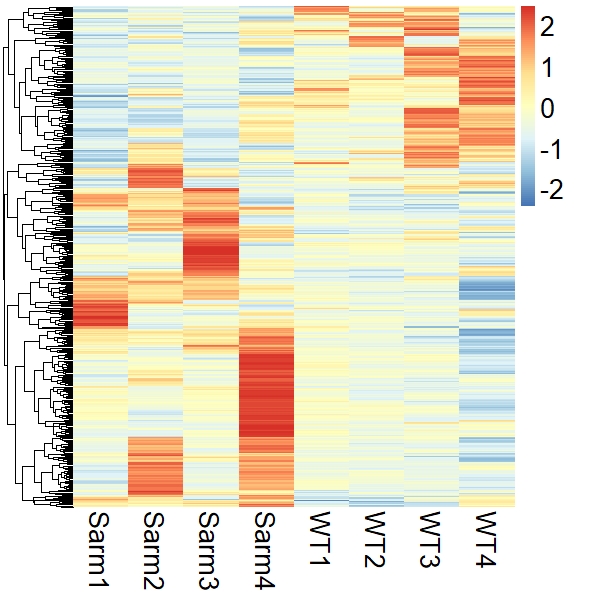

**Figure S2. Differentially expressed genes in surviving cortical neurons after stroke between WT and *sarm1*-/- mice.** Heatmap of normalized differentially expressed genes (FDR<0.05) in individual samples of MACS-FACS isolated cortical neurons after stroke (*n*=4/genotype). Color bar indicates log fold-change (logFC).

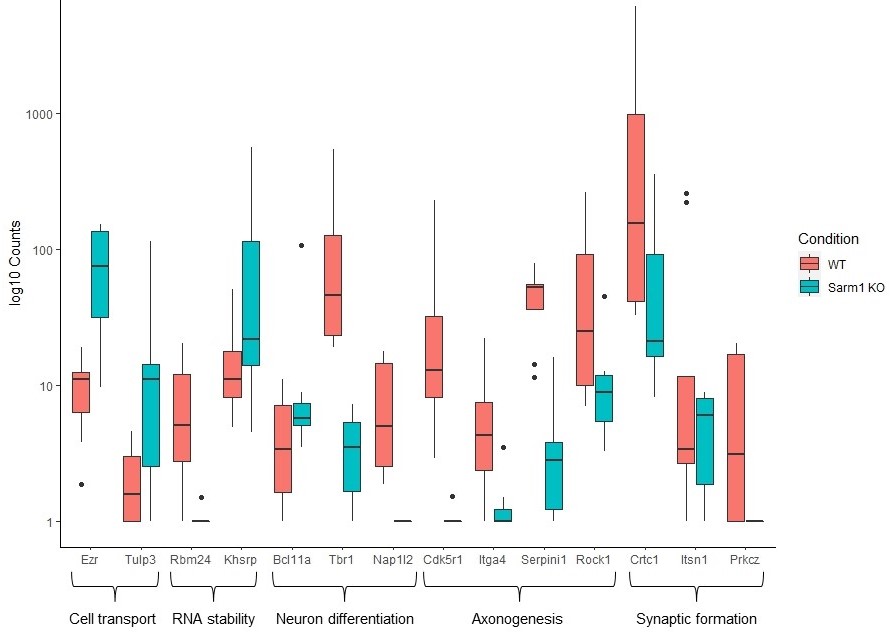

**Figure S3. Expression levels of hallmark genes from molecular program clusters in *sarm1*-/- stroke-injured cortical neurons.** Box plot of normalized read counts (log_10_) between wild-type (WT) and *Sarm1*-/- for selected differentially expressed genes (FDR<0.05) from each of the five major gene ontology clusters identified by REVIGO in MACS-FACS isolated stroke-injured cortical neurons.

**Axonogenesis Cluster	Synaptogenesis Cluster**

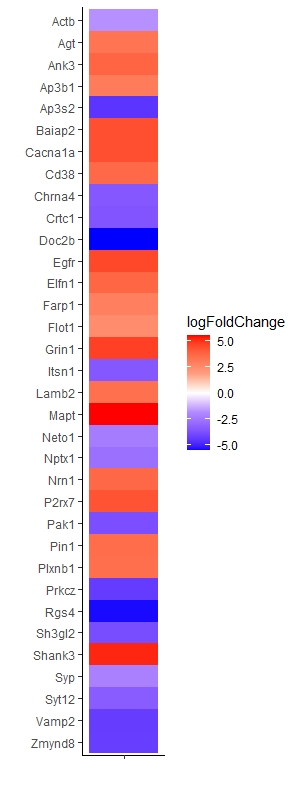

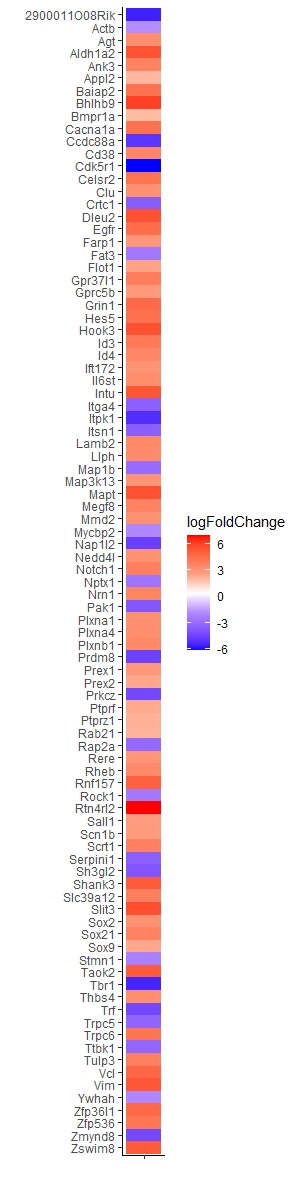

**Figure S4. Genes within the axonogenesis and synaptogenesis clusters.** Heatmaps of logFC (red=up; blue=down) for genes (all FDR<0.05) within the axonogenesis (left) and synaptogenesis (right) gene ontology clusters enriched in *Sarm1*-/- stroke-injured cortical neurons.

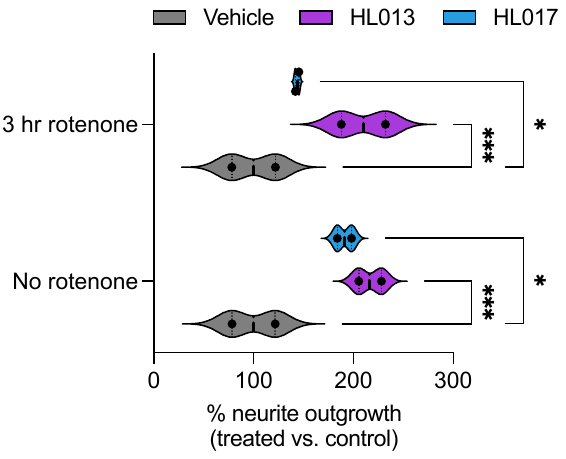

**Figure S5. Hit compounds from functional genomics screen drive neurite outgrowth after chemical ischemia.** Primary cortical neurons cultured for 7 days were treated with vehicle or rotenone (25 μM in 0.5% DMSO) for 3 hours, after which the cells are washed and replaced with fresh culture media. Treatment with rotenone resulted in an average 93.7% ± 0.46% reduction in absolute neurite length compared to no rotenone when measured 48hrs later (at day 9 *in vitro)*. Cultured cortical neurons exposed to vehicle or rotenone for 3 hours on day 7 *in vitro* were then immediately washed and further exposed to drug treatment with either vehicle, HL013 (purple), or HL017 (blue) for another 48 hrs. After drug treatment, neurite outgrowth was measured by calcein-AM on day 9 *in vitro* and normalized in all conditions to post-rotenone neurite length. In the no rotenone condition, drug treatment drives neurite outgrowth as previously shown (Fig. 4). In rotenone treated neurons, drug treatment also significantly promoted neurite outgrowth (HL013 vs. control – 209.9% ± 31.7% vs. 100% ± 31.0%, *p*=0.0006; HL017 vs. control – 143.6% ± 2.5% vs. 100% ± 31.0%; *p*=0.0161). Assays performed in technical duplicate.

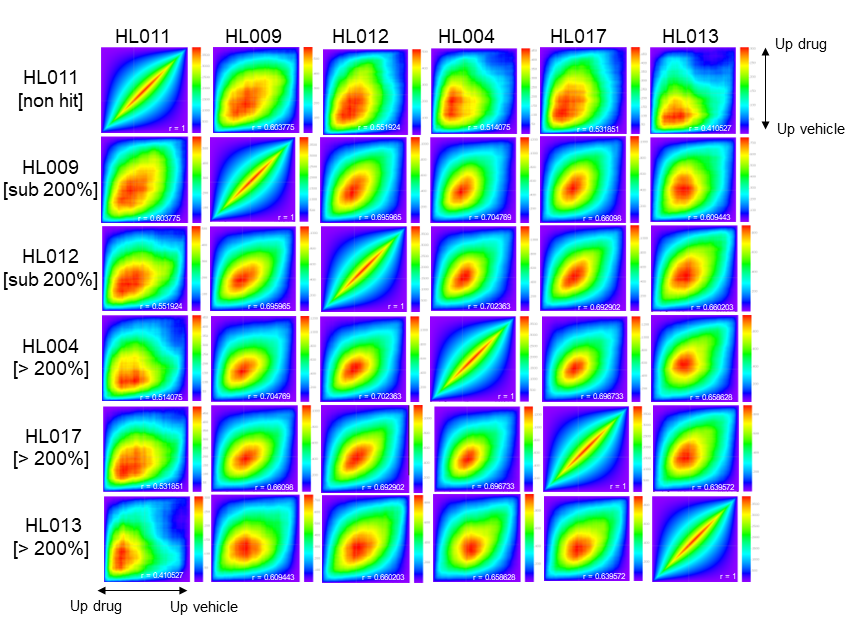

**Figure S6. Coordinated epigenetic signature of compounds identified by functional genomics screen.** Differentially accessible genes identified by ATAC-seq from hit compound treated cells were compared using the rank-rank hypergeometric overlap algorithm. These comparisons of gene accessibility profiles in neuron cultures treated with functional genomics hit compounds were all statistically significant (*p*<0.0001). Neuron cultures demonstrating significant neurite outgrowth 48 hours after treatment with hit compounds (HL004, HL009, HL012, HL013, HL017) demonstrated consistently higher hypergeometric overlaps (*r*>0.6) compared to those that did not demonstrate significant neurite outgrowth (HL011).

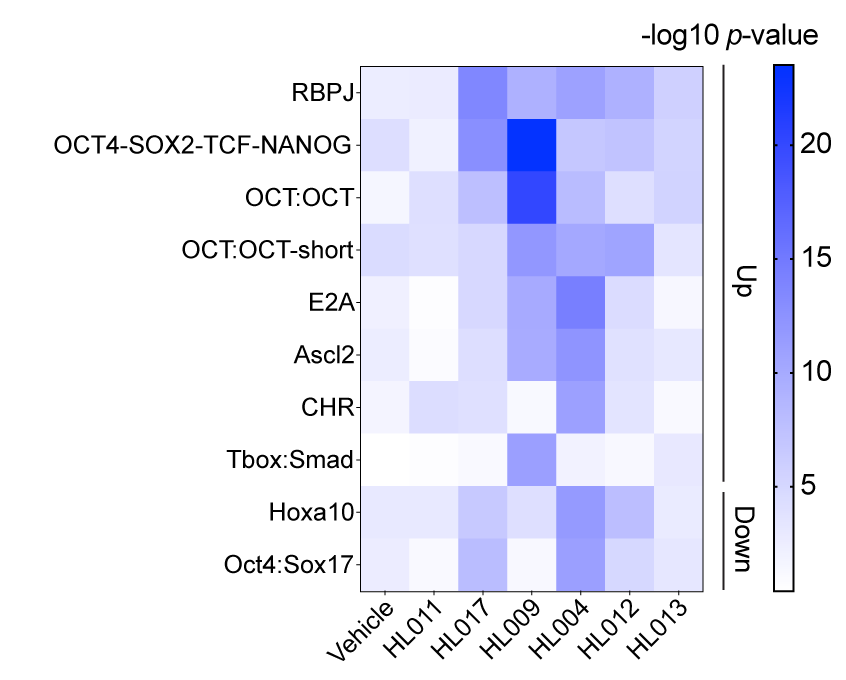

**Figure S7. Shared transcription factor motif analysis in neurite outgrowth promoting compounds.** Motif enrichment analysis by drug condition showing the -log10 *p*-value of transcription factor (TF) motif enrichment. TF motifs from ATAC-seq results were filtered for common up-regulated and down-regulated motifs across all drug conditions. Shared motifs were compared revealing significant enrichment of motifs associated with neurite and axon outgrowth in hit compound conditions compared to vehicle treated neurons. Consensus sequences for transcription factor motifs are available in SI Table 2, Sheet 3.

**Table S1. Loss of *Sarm1* up-regulates pro-growth genes in stroke-injured cortical neurons.**

| **Functional categories of top 150 up-regulated genes in *Sarm1*-/- stroke-injured cortical neurons** | | | | | | | |
| --- | --- | --- | --- | --- | --- | --- | --- |
| **Neuronal Differentiation** | | |  |  | | |  |
| *Mdga1* | *Lbh* | *Rtn4rl2* | *Cd46* | *Fbrs* | | | *Dact3* |
| *Pou3f1*  *Hook3* | *Heg1*  *H3f3a* | *Lurap1*  *Zswim8* | *Ccm2*  *Shcbp1l* | *Tbcb*  *Aldh1a2* | | | *Rnf157* |
| **RNA Processing** | | |  |  | | |  |
| *Wdtc1*  *Mettl3* | *Hint1* | *Rpl10a* | *Mbp* | *Rbmx* | | | *Dleu2* |
| **Synaptic Regulation** | | |  |  | | |  |
| *Syap1* | *Rock1* | *Sdk2* | *Intu* | *Bin2* | | | *Pcdhgc4* |
| **Cellular Transport** | | |  |  | | |  |
| *Sil1* | *Ap4e1* | *Hrg* | *Lin7b* | *Tacc1* | | | *Angpt2* |
| *Sptb* | *Slc6a20a* |  |  |  | | |  |
| **Ion Homeostasis** | | |  |  | | |  |
| *Slc30a3* | *Slc7a8* | *Slc20a2* | *Nkain2* | *Slc25a36* | | | *Atp7b* |
| *Atp1a4* | *Xcr1* | *Abcb7* | *Gclm* |  | | |  |
| **Neuronal Metabolism** | | |  |  | | |  |
| *Pyurf* | *Timp3* | *Ppp2r3c* | *Nmt1* | *Hs6st2* | | | *Umps* |
| *Cenpf* | *Sae1* | *Olfml2a* | *Adamts4* | *Gnai3* | | | *Pde5a* |
| *Bckdk* | *Zan* | *Glb1* | *Pou6f1* | *Ifnar1* | | | *Mtf2* |
| *Gstm5* | *Spata2* | *Fads2* | *Trmt61a* | *Atg3* | | | *Tdpoz2* |
| *Plcl1*  *Sucla2* | *Htra1*  *Cpa2* | *Mst1*  *Suox* | *Npepps* | *Pik3ca* | | | *Ube2r2* |
| **Cell Death Regulation** | | | | | | | |
| *Ltbr* | *Spop* | *Ercc5* | *Usp7* | |  |  | |
| **Axon outgrowth** | | | | | | | |
| *Lama5* | *Bhlhb9* | *Slit3* | *Mapt* | *Taok2* | | | *Shank3* |
| *Ngrn* | *Ccdc68* | *Ina* | *Vim* | *Krt25* | | |  |
| **Transcriptional Control** | | | | | | | |
| *Msantd1* | *Crat* | *Klf13* | *Mapk3* | *Prdm15* | | | *Myt1l* |
| **Unknown Function** | | |  |  | | |  |
| *Ccdc191* | *Stmn1-rs1* | *Brip1os* | *Fam13b* | *Tmem121b* | | | *Abracl* |
| *Mir6374* | *Tmem60* | *Gm2716* | *Krt90* | *Dnah2* | | | *Ints11* |
| *Ccdc127* | *Dhrs1* | *H2-Eb2* | *Gm20744* | *Zxdb* | | | *Ankrd24* |
| *Hcfc1r1* | *Plut* | *Utp11* | *Vmn2r57* | *Fbxw18* | | | *Tmem51* |

Unnamed genes within top 150 up-regulated genes are not included.

**Table S1. Functional categories of top 150 up-regulated genes in *sarm1*-/- stroke-injured cortical neurons.** Loss of *Sarm1* up-regulates pro-growth genes in stroke-injured cortical neurons. RNA-seq differential gene expression data from stroke-injured cortical neurons between *sarm1-/-* and wild-type mice (*n*=4/genotype) seven days after stroke**.** Unnamed genes within the top 150 up-regulated genes are not included.

**Table S2. Wild-type vs. *Sarm1-/-* Stroke-injured Neuronal Transcriptome.** RNA-seq differential gene expression data from stroke-injured cortical neurons between *Sarm1-/-* and wild-type mice (*n*=4/genotype) seven days after stroke including significant genes (FDR<0.05; Sheet 1), expressed genes (Sheet 2) and full DESeq output (Sheet 3).

**Table S3. Gene Ontology Analysis.** List of gene ontology terms generated from the differentially expressed *Sarm1-/-* stroke-injured cortical neuron transcriptome used as input for REVIGO query (Sheet 1). REVIGO output data including x,y semantic space coordinates and significance values (Sheet 2).

**Table S4. CMap Functional Genomics Screen Data.** CMap query results for compounds generated using the differential gene expression data from the top 150 genes (up- and down-regulated) and top 150 down-regulated genes. Compound names are listed along with combined CMap scores. HL library compounds were selected by further curation for known or predicted blood-brain permeability characteristics.

**Table S5. Drug-induced Differential Gene Accessibility Data.** ATAC-seq results from HL library drug exposures in E18 cortical neurons compared to vehicle controls (Sheet 1). Up-regulated gene ontology terms enriched in drug vs. vehicle comparisons (Sheet 2). Transcription factor motif consensus sequences enriched in drug vs. vehicle comparisons (Sheet 3).
